# Ecological resource competition as a driver of metallome evolution

**DOI:** 10.1101/2024.12.11.627971

**Authors:** Morgan S. Sobol, Holly R. Rucker, Eric Libby, Betül Kaçar, Ariel D. Anbar

**Affiliations:** Department of Bacteriology, University of Wisconsin-Madison, Madison, WI, USA; Department of Biology, Texas State University, San Marcos, TX, USA; Department of Mathematics and Mathematical Statistics, Umeå University, Umeå, Sweden; School of Earth and Space Exploration and School of Molecular Sciences, Arizona State University, AZ, Tempe, USA

**Keywords:** nitrogenase, molecular paleobiology, GOE, Proterozoic, molybdenum, metallome, resource competition

## Abstract

The oldest nitrogenase isozyme, emerging a billion years or more before the Great Oxidation Event (GOE), required a molybdenum (Mo)-based cofactor. “Alternative” nitrogenases using iron (Fe) or vanadium (V) cofactors evolved only after the GOE. This history is puzzling because environmental Fe availability *decreased* after the GOE, while Mo availability *increased*, due to the contrasting environmental redox behaviors of these elements. Why, then, did the alternatives emerge only after the GOE? Using a model constrained by known microbial Mo quotas, we demonstrate that a strong selection pressure for the use of metals in nitrogenase other than Mo is a plausible consequence of competition between nitrogen-fixing prokaryotes and nitrate-reducing microbes, which require Mo for nitrate reduction and assimilation. This competition would have intensified after the GOE due to increasing availability of nitrate, explaining the evolutionary timing of nitrogenase’s isozymes. Ecological resource competition therefore emerges as a third driver of metallome evolution in deep-time, alongside the relative environmental availabilities and adaptive advantages of particular metals.

## Introduction

Element use in biology is intricately linked to element availability in the surrounding environment. Consequently, metalloenzyme evolution and biological stoichiometry are broadly thought to have been shaped by changes in environmental availabilities throughout Earth’s history (1–5). However, the chemical characteristics of an element that make it biochemically useful also affect element selection, leading to selection pressures that can be in tension with availability (6). A third possible driver in the evolution of metal-containing biomolecules within a biological system (i.e., the metallome) is largely unexplored: resource competition, and its impact on organismal interaction and community composition (6).

The importance of resource competition may be seen in the evolution of biological nitrogen fixation and the enzyme “nitrogenase”, which arose by the mid-Archean (7–10). All life depends on this biological process, mediated by microorganisms called diazotrophs, and thus on nitrogenase, through which nitrogen enters the biosphere (∼270 – 310 Tg N year^-1^ (11)) by being “fixed” from N_2_ into a chemically reduced form that can be used to build new nucleic acids, proteins, and other biomolecules (12). Nitrogenase is a metalloenzyme existing in three distinct isoforms, containing either molybdenum and iron (“Mo-nitrogenase”), vanadium and iron (“V-nitrogenase”), or only iron (“Fe-nitrogenase”) (**Figure 1A** and **B**). Today, Mo-nitrogenase is the most abundant form; the Mo-independent nitrogenases are collectively termed “alternative” nitrogenases (13–15). Parsimony may suggest that the metal preferences of nitrogenases were shaped primarily by changes in environmental availability, whereby the biosphere optimized around the most abundant metal at its disposal at a given time on an evolving planet. If so, Fe-nitrogenase would be ancestral because Earth’s oceans at the time of nitrogenase’s emergence were O_2_-free and hence replete with Fe (16–18), while Mo was likely scarce (19). However, geochemical and evolutionary studies independently indicate that ancestral nitrogenases favored Mo (8, 20, 21) (**Figure 1C**). Phylogenetic reconstructions further suggest that alternative nitrogenases emerged through an ancient subfunctionalization of a Mo-nitrogenase at least 1 billion years later (9, 22) (**Figure 1A** and **1C**), and the acquisition of characteristic sequence novelties (i.e., the G subunit) (23) whose birth coincides with the approximate timing of the Great Oxidation Event (GOE) (10, 22–25).

**Figure 1.**
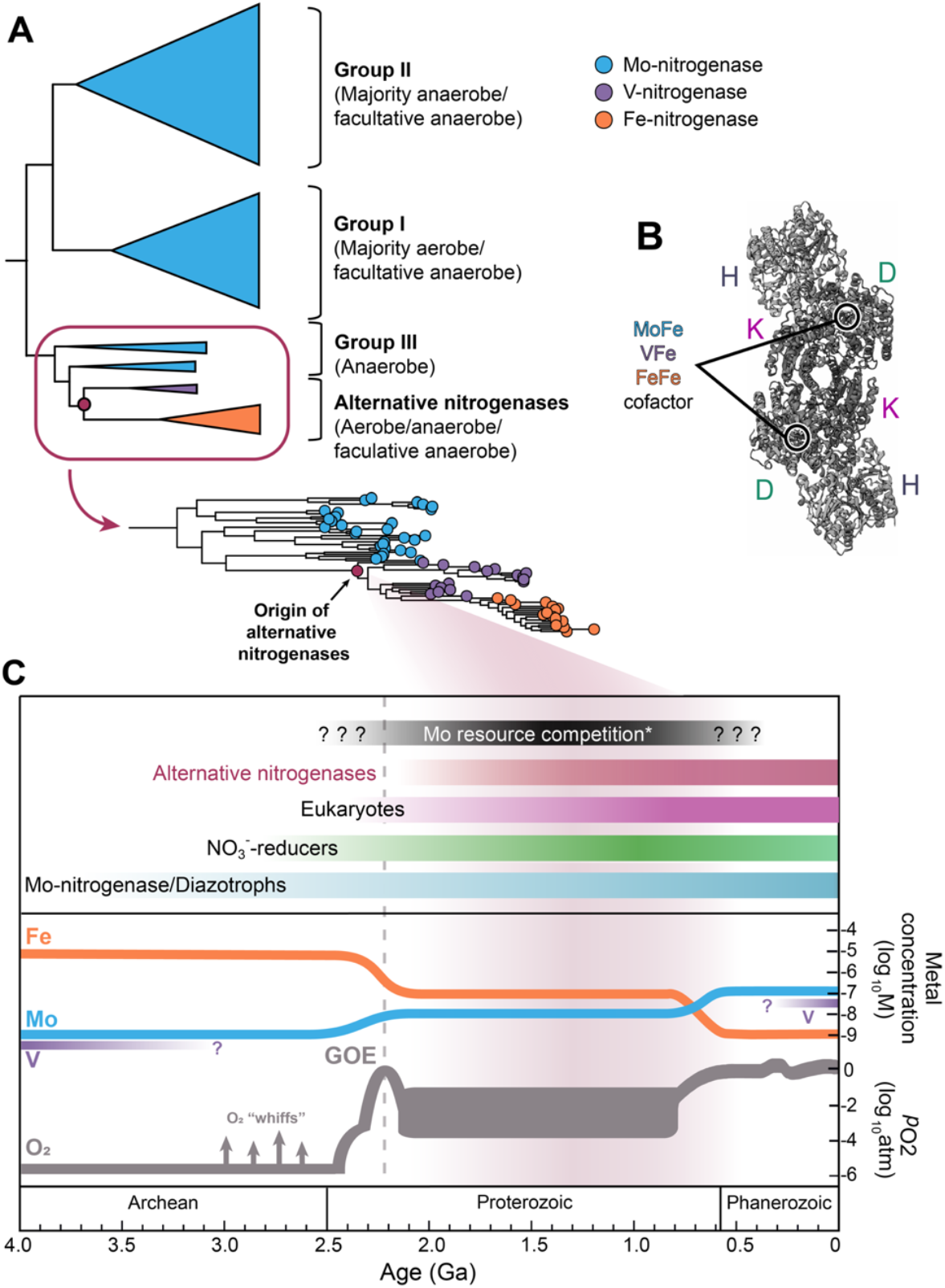
Key evolutionary events surrounding the evolution of alternative nitrogenases. **A)** Evolutionary history of nitrogenase based on maximum-likelihood phylogenetic construction of nitrogenase structural genes HDK (9) with aerobic, anaerobic, and facultative clades highlighted. Node distances are not to scale. **B)** Structure of the *Azotobacter vinelandii* Mo-nitrogenase complex (HDK) (57) and the three metal cofactors that different nitrogenase isozymes variably incorporate (Mo, V, and Fe) (58–60). **C)** Estimated age of alternative nitrogenases coincides with the Great Oxidation Event (GOE), and consequent changes in Mo, V, and Fe availabilities. The era of hypothesized competition for Mo between diazotrophs and ancestral NO_3_^-^-reducers (black bar) presumably began prior to the GOE, when evidence of ocean oxygenation appears (61), and persisted until pervasive environmental oxygenation led to a sharp rise in Mo ocean availability in the Neoproterozoic (62). Atmospheric O_2_ partial pressure estimates through time are from (63). Ocean metal abundances for Mo and Fe are from (64). Vanadium availability is estimated from (65, 66). Nitrogenase age estimates are based on geological nitrogen isotope signatures (8), evolutionary rate analysis (22), and molecular clock analysis (10, 24, 67). Time range for NO_3_^-^-reducers is based on molecular clock analysis (10) and geological nitrogen isotope signatures (8, 68). The time range for eukaryotes is based on molecular clock analyses and fossil evidence (51, 52, 54, 55).

The early emergence of Mo-nitrogenase despite Mo scarcity in Archean oceans may be understood in terms of its greater efficiency in N_2_ reduction, as it requires less ATP than its counterparts (14, 26, 27). However, the emergence of the Mo-free alternatives after the GOE remains paradoxical: Why would the less efficient Mo-free alternative nitrogenases evolve during a time when Mo availability *increased*, and Fe availability *decreased*? What evolutionary pressures drove the diversification and selection for alternative nitrogenase metal cofactors at that time?

We propose that the complex evolutionary history of the nitrogenases is best explained by ecological resource competition. Although our current understanding of resource competition offers valuable insights into modern microbial ecology (28, 29), its role is often overlooked in studies of early microbial life.

Today, the distribution of diazotrophs in global oceans is regulated by a complex balance among many factors, known and unknown. Known factors include nutrients, sunlight, predation, and competition with non-diazotrophs (30–33). Iron has been the focus of many of these studies due to its role as a limiting nutrient, given nitrogenase’s high Fe requirements (30, 34, 35). Prior work has shown that diazotrophs and NO_3_^-^-assimilators can co-exist if the ratio of Fe:N favors N_2_-fixation, as diazotrophs typically require more Fe than non-diazotrophs (34, 36), aligning with observations of N_2_fixation in NO_3_^-^ replete ocean regions (see (37); but also (38–42). In contrast, analogous ecological implications of Mo concentrations have not been explored in this framework, likely because Mo is not a limiting resource in modern oceans (43). Our current knowledge tells us that Precambrian oceans had 10-100x less Mo than in today’s oceans (44) (**Figure 1**), likely making it a limiting resource. This raises the question: If Mo was limiting in ancient oceans, could competition for it have contributed to the evolution of alternative nitrogenases (**Figure 2A**)?

**Figure 2.**
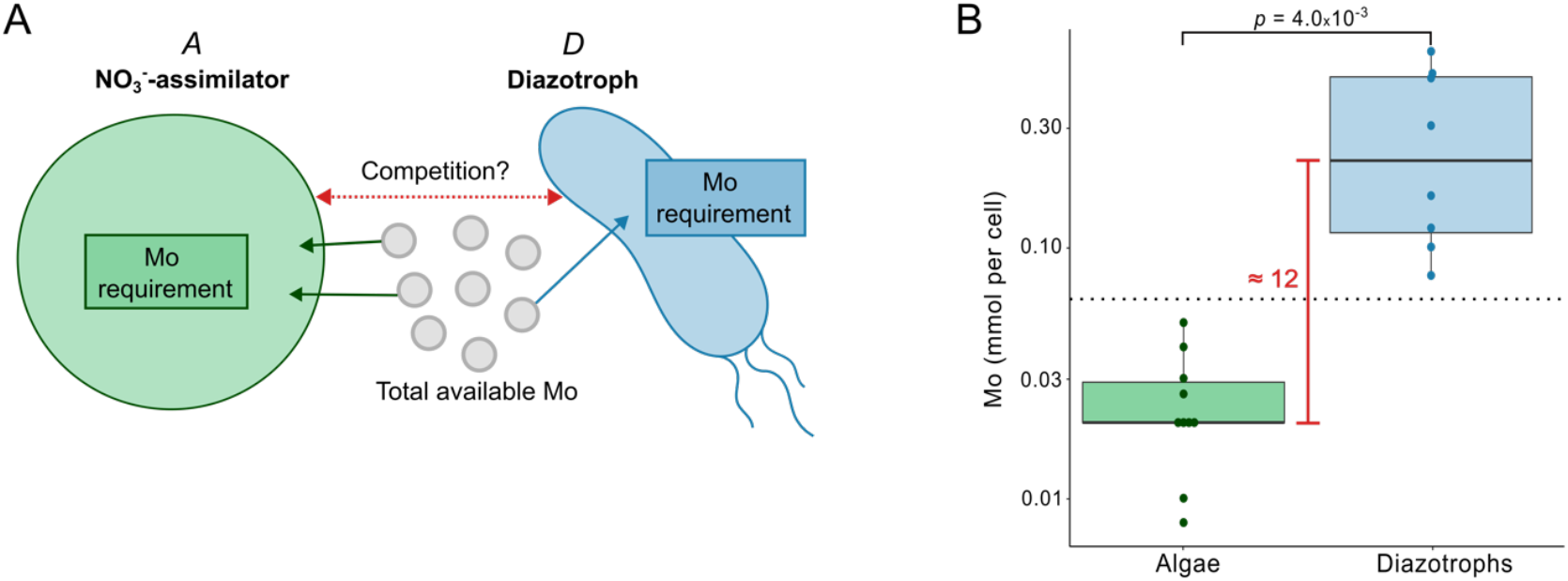
Basis of the competition hypothesis. **A)** Schematic of the hypothesized competition. **B)** Empirical estimates of Mo requirements from extant diazotrophs (blue) and algae (green), our model NO_3_^-^-assimilators, were compared. Diazotrophs require significantly more Mo than NO_3_^-^-assimilators on average. We estimated the median Mo requirement difference between diazotrophs and NO_3_^-^-assimilators to be 12 (red solid line). The black dotted line represents the midpoint between diazotrophs’ lowest Mo requirement and NO_3_^-^-assimilators’ highest. The y-axis is presented on a log_10_ scale to better visualize differences across orders of magnitude. Data can be found in Supplemental Table 2.

Our hypothesis builds on observations suggesting that nitrate (NO_3_^-^) concentrations in surface oceans increased after the GOE in part due to oxidative remineralization of biomass (45–48). The increasing availability of NO_3_^-^and Mo together favored the ecological radiation of NO_3_^-^-reducing microbes, including both prokaryotes and eukaryotes, which use another Mo-dependent enzyme: NO_3_^-^-reductase (49). Genes utilized in NO_3_^-^ reduction are inferred to have arisen around the time of the GOE (2.8-2.3 Ga) (10) (**Figure 1C**). We propose that competition over a Mo and NO_3_^-^supply that was rising, but still limiting to the demands of existing diazotrophs and emerging NO_3_^-^-reducers—which emerged between 2.8 – 1.0 Ga (10, 50–56)—could have generated a selection pressure that drove the evolution and subsequent diversification of Mo-independent alternative nitrogenases.

To evaluate our hypothesis quantitatively, we consider empirical differences in cellular concentrations of Mo between diazotrophs and NO_3_^-^-assimilators and assess how these differences impact the competition dynamics between diazotrophs and NO_3_^-^-reducing eukaryotes that assimilate NO_3_^-^.

## Results and Discussion

We first refer to studies that report measured cellular Mo concentrations in diazotrophs (69–72) and in algal NO_3_^-^-assimilators (72) (**Figure 2B; Supplemental Table 2**). Here, we focus on algae as the selected representative of NO_3_^-^-assimilators because extensive data are available on their Mo requirements (72), and because the emergence of eukaryotes as NO_3_^-^-assimilators would have significantly impacted diazotrophs by increasing the competition for Mo (**Figure 1**). Normalized cellular Mo concentrations (reported as mmol Mo cell^-1^) range from 0.077 to 0.6 for diazotrophs, whereas for algae they range from 0.008 to 0.05 **(Figure 2B; Supplemental Table 2**). Thus, diazotroph cells contain, on average, 12 times more Mo than algal cells, representing an order of magnitude difference (one-tail *p*-value=4.0 x 10^-3^ ; **Supplemental Table 4**). Empirical differences in Mo requirements between diazotrophs and NO_3_^-^-assimilators can likely be attributed to differences in the catalytic reaction rates of their respective molybdoenzymes (73, 74).

We next adapted a model influenced by Monod and Droop frameworks (75, 76), used widely in ecological modeling (77), to assess how the differences we observed in Mo requirements may lead to competition for Mo when Mo is the limiting resource (see **Supplemental Text** and **Supplemental Figure 1** for model details). Diazotrophs (*D*) and NO_3_^-^-assimilators (*A*) each have their own Mo requirements (mmol Mo cell^-1^) for growth (termed *θD* and *θA*, respectively). Using empirical estimates (**Figure 2B; Supplemental Table 2**), we focus the outcome of the competition on the theoretical uptake of Mo (*k*) that would be required for diazotrophs to survive. We use the term *k* to represent diazotrophs’ relative competitive effectiveness in capturing Mo for their private use (e.g., Mo transporter affinity, Mo internal storage, molybdophores), compared to NO_3_^-^-assimilators.

If maximal possible growth rates are the same for both species, the competition outcome is directly determined by their respective Mo requirements. If the median difference in Mo requirements is 12 (**Figure 2B**), diazotrophs must sequester at least 12 times more Mo than NO_3_^-^-assimilators in order to survive (**Figure 3**). The pressure on diazotrophs to efficiently sequester Mo can instead be alleviated if they are able to reduce their high Mo requirement (i.e. moving left along the x-axis in **Figure 3**). This logic applies under different growth rate scenarios, unless the net maximal growth rate of the diazotroph is 12 times greater than that of the NO_3_^-^-assimilator. It is not likely that diazotrophs would achieve such high rates when the two species cohabitate, as previous models and experimental work suggest that diazotroph growth rates are lower than or comparable to those of non-diazotrophic phytoplankton (31, 78–81). Other modeling approaches which are not dependent on growth rate differences can be used to uncover similar results in terms of the ratios of Mo requirements (see Supplemental Text for an example using the Droop equation). Thus, the central message is reinforced: diazotrophs could face severe competitive pressure due to their high Mo demands, regardless of growth rate differences, as long as an external supply of fixed N sustains NO_3_^-^-assimilator populations. Given this constraint, we find three evolutionary strategies that could emerge to help reduce the high Mo requirement of diazotrophs.

**Figure 3.**
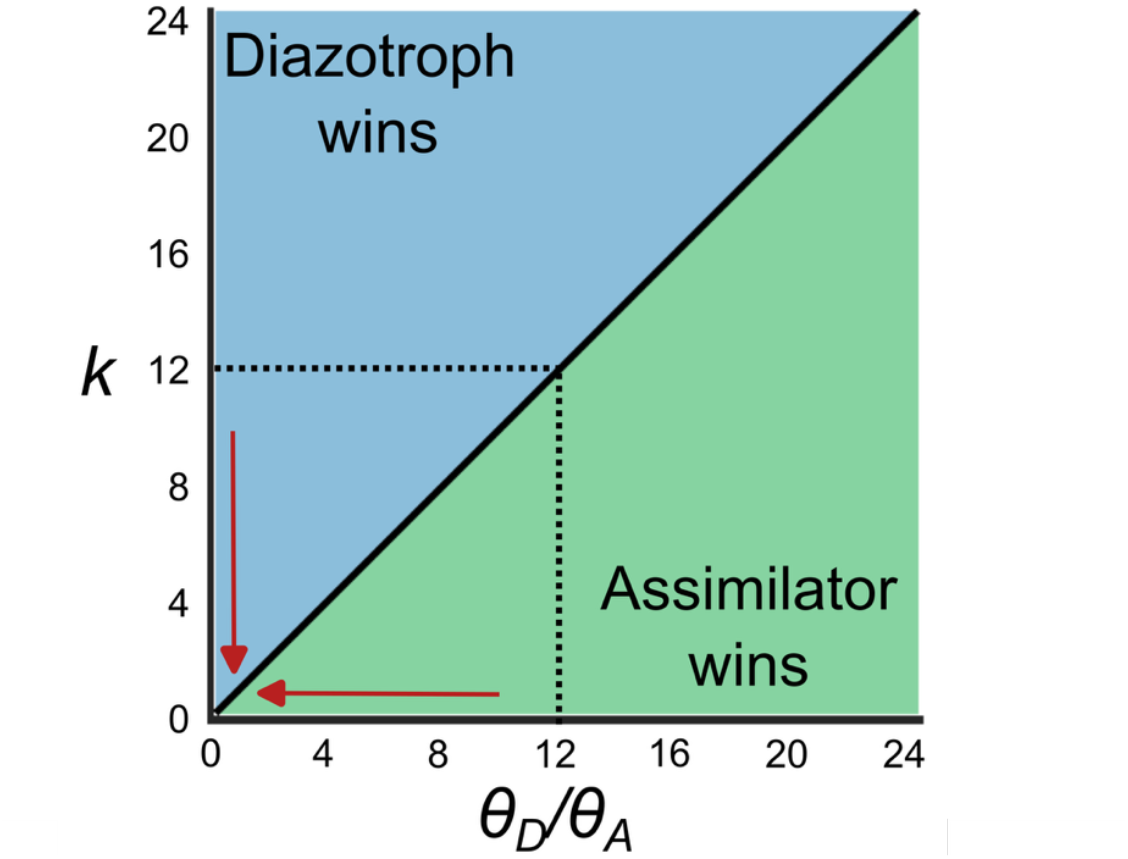
Mo competition model outcome. Competition dynamics based on differences in Mo requirements (*θ*_*D*_⁄*θ*_*A*_) and diazotrophs (*D*) relative competitive effectiveness in the uptake of Mo (*k*) compared to NO_3_^-^-assimilators (*A*). The estimated ratio of specific Mo requirements of diazotrophs relative to algae (*θ*_*D*_⁄*θ*_*A*_) is ∼ 12 (black dotted lines), as suggested by the empirical data assembled here. When the growth rates for NO_3_^-^-assimilators and diazotrophs are the same, the ratio of *θ*_*D*_⁄*θ*_*A*_ to *k* can be described as a proportional relationship that determines who wins the competition. In other words, when *θ*_*D*_⁄*θ*_*A*_ is equal to 12, then diazotrophs need to sequester at least 12x more Mo than NO_3_^-^-assimilators in order to survive. As the relative Mo requirement decreases (red arrows moving left on the x-axis), the “win condition” for diazotrophs can be achieved with smaller values of *k*, reducing the need for highly efficient Mo sequestration. As long as diazotrophs do not have a net maximal growth rate that is 12x greater than NO_3_^-^-assimilators, the same logic applies. The evolution of alternative nitrogenases is therefore a probable strategy to cope with competition for Mo.

First, diazotrophs could shift to the assimilation of NO_3_^-^, effectively lowering their Mo cost to levels comparable to NO_3_^-^-assimilators. However, if fixed N is limited, this strategy may lead to direct competition for both NO_3_^-^ and Mo.

Alternatively, diazotrophs could evolve mechanisms to capture and store Mo more efficiently than NO_3_^-^-assimilators. However, whether diazotrophs can achieve this relative level of sequestration efficiency is not yet well understood. Among the few model diazotrophs examined, reported Mo uptake rates vary widely from *K*_*m*_ ≈ 0.3 nM in *Anabaena* to >50 nM in root-nodulating *Bradyrhizobium japonicum* (82, 83). But, field observations from Lake Cadagno, a Proterozoic Ocean analog, have implicated that purple sulfur bacterial diazotrophs encode high-affinity Mo transporters (*K*_*m*_yet to be determined) that enable N_2_fixation in this low Mo environment (84). Comparable data for algae is limited, but two molybdate transporters from *Chlamydomonas reinhardtii*, MOT1 and MOT2, were reported with *K*_*m*_values ≈ 6 nM and ∼550 nM, respectively (85, 86). Mo storage has not been reported in algae and does not appear to be a widespread mechanism among diazotroph populations (70, 87), nor does the advantageous, albeit expensive, production of extracellular Mo-scavenging metallophores (88, 89).

Finally, the constraints imposed by Mo competition favor a third strategy: the evolution of Mo-free, alternative nitrogenases. By shifting to a Mo-independent pathway for N_2_-fixation, diazotrophs can bypass the selection pressure for Mo experienced when cohabiting with NO_3_^-^-assimilators under N and Mo limitation. Thus, the evolution of alternative nitrogenases offers a superior solution by decoupling N_2_-fixation from Mo availability altogether.

This competition dynamic becomes especially relevant when considering the evolutionary pressures faced by diazotrophs in ancient Mo-limiting environments. As NO_3_^-^ availability in the oceans increased during the Proterozoic, fueling the emergence and ecological advancement of Mo-using NO_3_^-^-reducers, competition for Mo likely intensified, disadvantaging diazotrophs that depended solely on Mo-nitrogenase and hence favoring the evolution of alternative nitrogenases.

The ability to extend our analysis to all NO_3_^-^ reducers is limited by the dearth of comparable Mo data for both dissimilatory and assimilatory NO_3_^-^ reducers. The Mo requirements for different NO_3_^-^ reductases are suggested to be influenced by variations in enzyme structure and active site configuration (90). However, the extent to which these properties impact overall cellular Mo requirements is poorly understood. Investigating whether the Mo requirements of NO_3_^-^-assimilators can be broadly extrapolated to NO_3_^-^ reducers should be a focus for future research. Similarly, the lack of Mo data for most diazotrophs limits our understanding of how Mo requirements may vary, underscoring the need for further studies. Addressing these gaps would also benefit from understanding Mo uptake dynamics between NO_3_^-^-reducers and diazotrophs in modern systems where Mo is limiting. Lastly, future work could test multi-variable frameworks that implement co-limitation of Mo alongside other nutrients (e.g. NO_3_^-^, Fe, P) and resources (e.g. light, temperature) to provide a more comprehensive view of the complexities that influence the interactions between diazotrophs and non-diazotrophs (34, 91–93).

Nonetheless, it appears that an ancient need for Mo left a lasting imprint at the molecular level, where alternative nitrogenases and Mo-sequestration strategies persist today (43, 67, 82, 84, 85, 87, 94–96). Most likely, alternative nitrogenases continue to provide a competitive advantage, functioning as “backup” enzymes in Mo-limited environments.

This study focuses on the consequences of ecological resource competition for the evolutionary origins of nitrogenase isozymes. Nitrogenase evolution is unique only in that we now have considerable knowledge about the timing of the evolution of metal usage in this enzyme family, alongside approximate contemporaneous changes in global ecology and the environmental availability of Mo. It is likely, however, that resource dynamics, including ecological resource competition as highlighted here, also played a significant role in shaping the evolution of other metalloenzymes, as resource competition is a fundamental driver of microbial interactions and evolution (29). Much remains to be uncovered about the integrated molecular, ecological, and geochemical histories of the metallome (6). Addressing this knowledge gap will be essential to advance our understanding of the co-evolution of Earth and life.

## Methods

We obtained normalized cellular Mo:P data from the literature of cultures grown with approximately 100 nmol L^-1^ of Mo for comparison (**Supplemental Table 2**). P is often measured simultaneously with metals (97), and so measured P data were used in these cases. When only C-normalized Mo quotas were provided, P-normalized quotas were calculated (**Supplemental Table 3**) using the Redfield ratio (C_106_N_16_P) (98). Statistical differences between diazotroph and algae Mo requirements were determined using the two-sample *t*-test assuming unequal variances with an alpha value of 0.05 (**Supplemental Table 4**).

## Acknowledgments

This work was supported by the NASA Interdisciplinary Consortium for Astrobiology Research: Metal Utilization and Selection Across Eons, MUSE [80NSSC17K0296] with additional support from the NASA Exobiology Program [NNH23ZDA001N] and the USDA-NIFA Hatch Grant [AWD00001400]. M.S. was supported by the NASA Postdoctoral Program, administered by Oak Ridge Associated Universities under contract with NASA. We thank Yang Kuang for extensive conversations during the development of an earlier version of this manuscript. We also thank Evrim Fer, Kaustubh Amritkar, members of the MUSE ICAR team, and earlier reviewers for their valuable feedback.

## Statement of authorship

conceptualization - AA, BK; designed research - MS, BK, AA; performed research - MS, HR; designed model - EL, MS; analyzed results - MS, HR, EL, BK, AA; wrote the paper - MS first draft; editing - all authors.

## Data accessibility statement

All data used in this study were obtained from previously published literature through the original sources cited in the manuscript.

