## Supplemental Information for "Ecological resource competition as a driver of metallome evolution"

### Supplemental Text

#### Original model

We build our model from existing premises that diazotrophs and $NO_3^-$-assimilators coexist when external N is limiting and Fe:N is favorable for $N_2$-fixation (1). Under this condition of low N concentration, diazotrophs that are capable of assimilating $NO_3^-$ will prefer to acquire N via $N_2$-fixation so as not to compete for $NO_3^-$ (2, 3). Thus, we consider a simple system in which populations (expressed as the number of cells) of diazotrophs ($D$) and $NO_3^-$-assimilators ($A$) cohabitate and compete for Mo (**Supplemental Figure 1**). Here, we focus on algae as the selected representative of $NO_3^-$-assimilators because extensive data are available on their Mo requirements (4), and because the emergence of eukaryotes as $NO_3^-$-assimilators would have significantly impacted diazotrophs by increasing the competition for Mo.

The modeled system (**Supplemental Figure 1, Supplemental Table 1**) considers that the total Mo pool, or $M_{total}$ (mmol Mo), from which both diazotrophs and $NO_3^-$-assimilators draw is fixed at some constant, but limited amount. We assume that diazotrophs and $NO_3^-$-assimilators each have their own Mo requirements (mmol Mo cell$^{-1}$) for growth ($\theta_D$ and $\theta_A$, respectively). Maximal growth rates (d$^{-1}$), $r_D$ or $r_A$, represent the theoretical highest growth rate that either species can achieve under ideal conditions. They serve as a benchmark, reflecting the potential for growth if Mo requirements ($\theta_D$ and $\theta_A$) are fully met. Population turnover $\phi$ per day (d$^{-1}$), or a population's removal from the environment, is assumed to be the same for both species, following traditional simplified resource-competition models (5–8). The allocation of Mo between the two species is therefore represented as a competition, whereby the total Mo pool is shared between the two species based on their population sizes and diazotrophs' relative competitive effectiveness in the uptake of Mo ($k$) for their private use, compared to $NO_3^-$-assimilators. Thus, as the populations of $NO_3^-$-assimilators and diazotrophs change over time, the amount of Mo (mmol Mo) used by each species ($M_D$ or $M_A$) also changes, altering their realized growth rates and subsequent survival. We use the form of a Monod equation $\frac{r_{cell}M_{cell}}{M_{cell}+\theta}$ (9) to describe the realized growth rates of diazotroph and $NO_3^-$-assimilator populations as a function of the amount of Mo per cell, or $Q_D$ and $Q_A$ (mmol Mo cell$^{-1}$).

To determine $M_D$ or $M_A$, we partition $M_{total}$ across the two species' populations depending on their relative abundances and the competition parameter *k*. Thus, the total amount of Mo used by diazotrophs ($M_D$) and $NO_3^-$-assimilators ($M_A$) can be described as:

*Equation 1:*

$$M_D = M_{total}\left(\frac{k\,D}{k\,D+A}\right)$$

*Equation 2:*

$$M_A = M_{total}\left(\frac{A}{k\,D+A}\right)$$

When $k = 1$, both species are equally matched in their ability to sequester Mo and so the total amount of Mo is distributed based on the fraction of each species in the population. Alternatively, when $k > 1$, diazotrophs can sequester more of the Mo supply. The inverse is true when $k < 1$. The amount of Mo (mmol) per cell (*Q*) is then the amount sequestered by diazotrophs ($M_D$) or algae ($M_A$) divided by the total population of each species (*D* or *A*), such that $Q_D = M_D/D$ and $Q_A = M_A/A$. Based on these considerations, we describe the population dynamics of *D* and *A* over time using a system of differential equations:

*Equation 3:*

$$\frac{dD}{dt} = r_D \frac{Q_D}{Q_D + \theta_D} D - \phi D$$

*Equation 4:*

$$\frac{dA}{dt} = r_A \frac{Q_A}{Q_A + \theta_A} A - \phi A$$

Within this system, the absence of one species allows the other species to grow to its carrying capacity, described by $\frac{(r_D - \phi) M_{total}}{\phi \theta_D}$ for *D* or $\frac{(r_A - \phi) M_{total}}{\phi \theta_A}$ for *A*. The carrying capacity for each species is determined when their growth is at equilibrium ($\frac{dD}{dt} = 0$ for *D* and $\frac{dA}{dt} = 0$ for *A*). When both species are present, competition for Mo results in one species driving the other to extinction, except for a very specific combination of parameters whereby the two species are equally competitive:

*Equation 5 - competition:*

$$k = \left(\frac{\theta_D}{\theta_A}\right)\frac{(r_A - \phi)}{(r_D - \phi)}$$

If $k > \left(\frac{\theta_D}{\theta_A}\right)\frac{(r_A - \phi)}{(r_D - \phi)}$, diazotrophs drive $NO_3^-$-assimilators to extinction, whereas if $k < \left(\frac{\theta_D}{\theta_A}\right)\frac{(r_A - \phi)}{(r_D - \phi)}$, then diazotrophs go extinct. The outcome of the competition, therefore, depends on three parameters: 1) the ratio of dependencies on Mo ($\frac{\theta_D}{\theta_A}$); 2) the ratio of maximal growth rates $\frac{(r_A - \phi)}{(r_D - \phi)}$; and 3) *k*.

To determine the ratio of Mo requirements ($\theta_D/\theta_A$), we refer to prior studies that reported measured cellular Mo concentrations in diazotrophs (4, 10–12) and in algal $NO_3^-$-assimilators (4) grown with approximately the same Mo concentration (~100 nmol $L^{-1}$) (**Supplemental Table 2**).

The results of our analyses of the original model can be verified numerically using the following code (below). The first is a MATLAB script called "run_original_model.m" that will run the competition and generate plots of the populations for *D* and *A* in time. It calls the second file called "compete_species_original_model.m" which is a MATLAB function, that encodes the dynamical system. By varying the choice of *k*, the outcome of the competition between *D* and *A* changes.

**run_original_model.m**

```
%% Run original model
% create a structure for the parameters (below is an example)
pars=struct();
pars.rD=.01;
pars.rA=.01;
pars.M=1000;
pars.thetaD=1;
pars.thetaA=0.5;
pars.phi=.001;

% critical k for coexistence is computed via equation 5 in the SI
critical_k=(pars.thetaD/pars.thetaA)*(pars.rA-pars.phi)/(pars.rD-pars.phi);

% k choices, its relationship to critical_k determines the outcome
% if k<critical_k then D goes extinct
pars.k=critical_k*(.9); % example
% if k>critical_k then A goes extinct
pars.k=critical_k*(1.1); % example
% if k=critical_k then A and D coexist
pars.k=critical_k; % example

% pick k
pars.k=critical_k*(1);
```

```
% running the competition, by solving the ODEs numerically using ode45
time_span=[0 100000]; % a time interval for ode45
initial_populations=[1000;1000]; % the initial D and A populations,
respectively
[t,DAsolution]=ode45(@(t,x)
compete_species_original_model(t,x,pars),time_span,initial_populations);

% extract the solution from the output of ode45
D=DAsolution(:,1);
A=DAsolution(:,2);

% plot the output
close all
plot(t,D,'LineWidth',3);
hold on;
plot(t,A,'LineWidth',3);
xlabel('Time','FontSize',24);
ylabel('Species','FontSize',24);
set(gca,'LineWidth',3,'FontSize',14)

% note if k=critical_k, the lines for D and A will overlap if their initial
% population sizes are the same.
```

**compete_species_original_model.m**

```
function [dvect]=compete_species_original_model(t,vect,pars)
% This function takes in three inputs and returns the derivative based on a
dynamical system of ODEs
% Inputs:
%  t: a real number representing "time". It has no effect on the output
since the ODEs are autonomous.
%  vect: a 2 X 1 vector of the populations of D and A respectively.
%  pars: a structure with the parameter values used in the ODE
% Outputs:
%  dvect: a 2 X 1 vector of the derivatives of the populations of D and A
respectively

% Assign the vector values to D and A concentrations
D=vect(1);
A=vect(2);

% Define some new terms corresponding to simplify the modeling
mD=pars.M*(pars.k*D)/(pars.k*D+A); % equation 1 in SI
mA=pars.M*(A/(pars.k*D+A)); % equation 2 in SI
qD=mD/D;
qA=mA/A;

% Derivatives for the dynamical system
dDdt=pars.rD*(qD/(qD+pars.thetaD))*D-pars.phi*D; % equation 3 in SI
dAdt=pars.rA*(qA/(qA+pars.thetaA))*A-pars.phi*A; % equation 4 in SI

% Format output
dvect=[dDdt;dAdt];
```

### Droop model

We consider a Droop model (13) that describes the growth of a species population determined by a limiting resource. Unlike the main model of the paper, the Droop model partitions Mo into internal and external stores and relates population growth to the amount of a nutrient in the internal stores. For example, the resulting set of equations for a single species (in this case $D$) is as follows:

*Equation 6:*

$$\frac{dD}{dt}=\mu_D(Q_D)D-\phi D$$

*Equation 7:*

$$\mu_D=max\left(r_D\left(1-\frac{\theta_D}{Q_D}\right),0\right)$$

*Equation 8:*

$$\frac{dQ_D}{dt}=p_D(M_{total})-\mu_D(Q_D)Q_D$$

*Equation 9:*

$$p_D(M_{total})=p_{max}\left(\frac{M_{total}}{K_{S,D}+M_{total}}\right)$$

*Equation 10:*

$$\frac{dM_{total}}{dt}=-p_D(M_{total})D+\phi Q_D D$$

The 6$^{th}$ equation describes the growth of the population of the species, where $D$ is the species concentration, $\mu_D$ is the growth rate of the species, which is a function of the internal concentration of Mo ($Q_D$), and $\phi$ is the turnover rate for the species.

The 7$^{th}$ equation describes the actual shape of the growth function ($\mu_D(Q_D)$). It is the maximum value of two terms, and so it varies between 0 and a maximum growth rate, $r_D$. The parameter $\theta_D$ describes the minimum amount of Mo needed for growth. If the internal concentration of Mo is above this threshold, i.e. $Q_D > \theta_D$, then the term $(1-\theta_D/Q_D)$ is greater than zero which means that the population increases. If instead $Q_D < \theta_D$ then the term $(1-\theta_D/Q_D)$ is negative so the population growth rate is set to zero, reflecting the fact that the population stops growing because of a lack of Mo.

The 8$^{th}$ equation describes the internal concentration of Mo per cell. The function $p_D(M_{total})$ describes uptake of Mo from the external environment. The resource uptake is a function of $M_{total}$, which is the Mo concentration outside the cell and the species rate of Mo uptake ($p_D$). Mo is therefore lost from the environment at a rate proportional to the species growth.

The 9$^{th}$ equation describes the resource uptake function. It is a sigmoid function in terms of the external Mo concentration $M_{total}$. The uptake function varies between 0 and a maximum rate $p_{max,D}$. The maximum rate is realized when $M_{total}$ is high, indicating that at this point the uptake rate is limited by the cell's transport capacity. The term $K_{S,D}$ is a parameter that determines when the uptake reaches half its maximum, i.e. $p_D(K_{S,D})=1/2\, p_{max,D}$.

The 10th and final equation for a species describes the dynamics of the external resource concentration. It decreases as cells take up Mo at a rate $p_D(M_{total})D$, which is the uptake rate per cell ($p_D(M_{total})$) multiplied by the number of cells ($D$). The external Mo concentration can also increase as cells die and release their contents back into the environment. This is captured by the term $\phi Q_D D$, which is the production of the turnover rate, the concentration of Mo per cell, and the concentration of cells.

This is the basic Droop model for growth of a single species. We can model a competition between two species by adding a parallel set of equations corresponding to the dynamics of the second species, $A$, and its internal Mo concentration $Q_A$. The one equation that directly contains terms for the competition between the two species is the one for $\frac{dM_{total}}{dt}$. The full generalized system for the two species is below:

*Equations for species D:*

$$\frac{dD}{dt}=\mu_D(Q_D)D-\phi D$$

$$\mu_D=max\left(r_D\left(1-\frac{\theta_D}{Q_D}\right),0\right)$$

$$\frac{dQ_D}{dt}=p_D(M_{total})-\mu_D(Q_D)Q_D$$
$$p_D(M_{total})=p_{max,D}\left(\frac{M_{total}}{K_{S,D}+M_{total}}\right)$$

*Equations for species A:*

$$\frac{dA}{dt}=\mu_A(Q_A)A-\phi A$$

$$\mu_A=max\left(r_A\left(1-\frac{\theta_A}{Q_A}\right),0\right)$$

$$\frac{dQ_A}{dt}=p_A(M_{total})-\mu_A(Q_A)Q_A$$
$$p_A(M_{total})=p_{max,A}\left(\frac{M_{total}}{K_{S,A}+M_{total}}\right)$$

*Equation 11 - competition:*

$$\frac{dM_{total}}{dt}=-p_D(M_{total})D+\phi Q_D D-p_A(M_{total})A+\phi Q_A A$$

We can analyze the outcome of the competition by assuming that one species is present (say $D$) and asking under what conditions $A$ can invade. We assume $D$ reaches a steady state so that $\frac{dM_{total}}{dt} = 0$, $\frac{dD}{dt} = 0$, and $\frac{dQ_D}{dt} = 0$. The steady state value of $Q_D$ (which we call $Q_D^*$) is $Q_D^* = \frac{r_D\theta_D}{r_D-\phi}$ and the steady state value of $M_{total}$ (which we call $M_{total}^*$) is: $M_{total}^* = \frac{K_{S,D}\,\mu_D(Q_D^*)Q_D^*}{p_{max,D}-\mu_1(Q_D^*)Q_D^*} = \frac{K_{S,D}\phi\, r_D Q_D}{p_{max,D}(r_D-\phi)-\phi\, r_D Q_D} = \frac{\phi\, K_{S,D}}{\left(p_{max,D}\right)\left(\frac{r_D-\phi}{r_D\theta_D}\right)-\phi}$.

In order for $A$ to invade, it must be such that $\frac{dA}{dt} \geq 0$. This implies that $\mu_A(Q_A) \geq \phi$. This means that $Q_A \geq \frac{r_A\theta_A}{r_A-\phi}$. If $A$ invades, then it must consume Mo from the external environment to cause $\frac{dM_{total}}{dt} < 0$. This means that $-p_A\left(M_{total}^*\right)+\phi Q_A < 0$. We can simplify by defining the ratio of uptake functions as some factor called $f$, which represents the relative competitive effectiveness of species $A$ in Mo uptake compared to species $D$, i.e. $f = p_A\left(M_{total}^*\right)/p_D\left(\mathrm{M}_{total}^*\right)$. Note if they are the same then $f = 1$.

We can use these relationships to find that species $A$ invades when $\phi Q_A < f\, p_D\left(M_{total}^*\right)$. We know $p_D\left(M_{total}^*\right) = \phi Q_D^*$ which we can plug in and rearrange terms to get $\left(\frac{r_D-\phi}{r_D}\right)\left(\frac{r_A}{r_A-\phi}\right)\frac{\theta_A}{\theta_D} < f$. If the turnover rate is low relative to the maximum growth rates then this simplifies to $\frac{\theta_A}{\theta_D} < f$.

We can now expand out $f$ to find: $\frac{\theta_A}{\theta_D} < \left(\frac{p_{max,A}}{p_{max,D}}\right)\left(\frac{K_{S,D}+M_{total}^*}{K_{S,A}+M_{total}^*}\right)$. If $\phi$ is small, then the value of $M_{total}^*$ is also small in comparison with the $K_{S,D}$ and $K_{S,A}$ terms. We can then simplify the condition for invasion to the equation below. The actual maximal growth rates do not factor into this competition so long as they are big enough for our simplifications.

*Equation 12 - invasion by A:*

$$\frac{\theta_A}{\theta_D} < \left(\frac{p_{max,A}}{p_{max,D}}\right)\left(\frac{K_{S,D}}{K_{S,A}}\right)$$

If this is satisfied, then species $A$ can invade.

If the uptake functions are the same, then this means that $A$ can only invade if it has a lower minimum threshold. Hence, if we set $\theta_A = 1$ and $\theta_D = 12$, reflecting our empirical estimates of minimum Mo cell quotas (**Supplemental Table 2**), the Equation for invasion of $A$ is satisfied. The result is that $A$ invades and replaces $D$.

The uptake functions can also determine the possibility of invasion. In order for $D$ to not be invaded by $A$, their overall uptake functions need to be greater. If, for example, $p_{max,A} = 1$ and $K_{S,A} = 0.5$, then $D$'s uptake function need to at least be $p_{max,D} = 2.4$ and $K_{S,A} = 0.1$ to not be invaded.

Finally, we note that changes to the turnover rate do not alter the results. If we use the same thresholds as before, i.e. $\theta_A = 1$ and $\theta_D = 12$ and uptake functions equal to 1, except we raise the turnover rate from 0.01 to 0.075, $A$ still invades.

Thus, we come to the same conclusion as in the original model: diazotrophs faced severe competitive pressure due to their high Mo demands, but could have compensated by either having a significantly higher uptake of Mo compared to $NO_3^-$-assimilators or by removing their dependence on Mo altogether via the use of alternative nitrogenases.

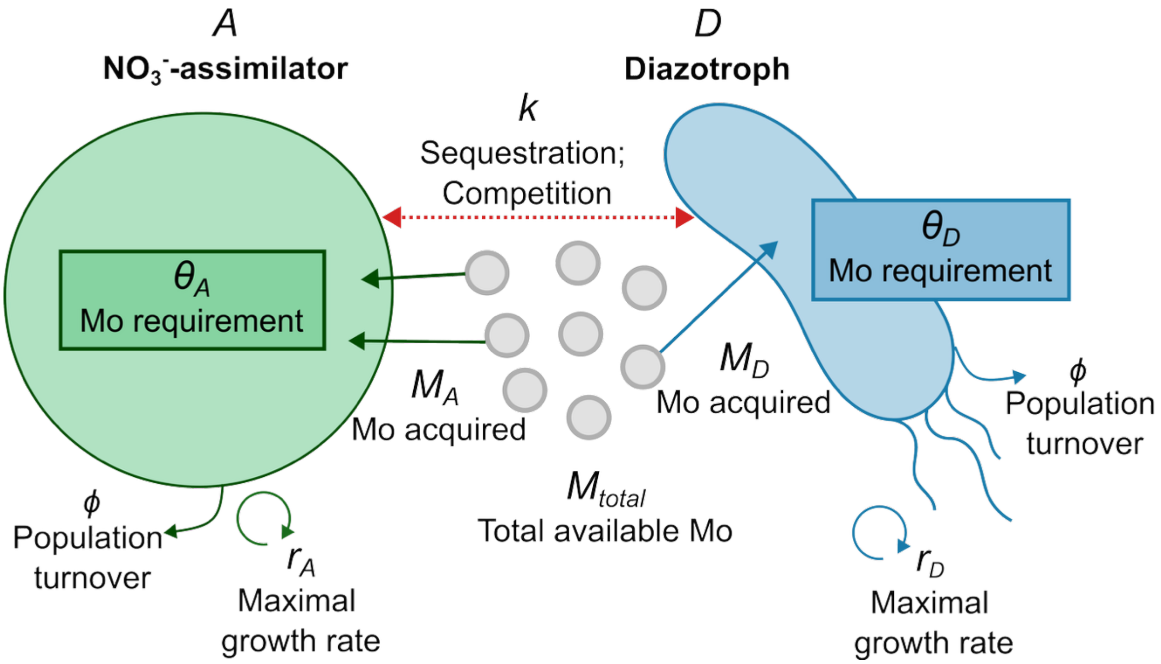


**Supplemental Figure 1. Overall components of the main proposed competition model.** Mo is required by both $NO_3^-$-assimilators and $N_2$-fixing prokaryotes (diazotrophs), thus in our model Mo (*M*) is the limiting factor for growth of the populations (expressed as the number of cells), of both $NO_3^-$-assimilators (*A*) and diazotrophs (*D*). Both species have their respective requirements for Mo (mmol Mo cell$^{-1}$), $\theta_A$ or $\theta_D$, and a maximal potential growth rate ($r_A$ or $r_D$), expressed as per day (d$^{-1}$). To determine the amount of Mo acquired by each species ($M_A$ or $M_D$), total available Mo, or $M_{total}$ (mmol Mo), is partitioned across the populations based on each organism's relative abundance and diazotrophs' relative competitive ability to uptake Mo (parameter *k*). Phi ($\phi$) is the loss rate parameter to account for population turnover per day (d$^{-1}$). Competition dynamics between $NO_3^-$-assimilators and diazotrophs can therefore be analyzed in terms of differences in their Mo dependency, maximal growth rates, and their ability to sequester Mo.

**Table S1.** Model variables and meanings

| Main model variables | Droop model variables | Variable definition | Units |
|---|---|---|---|
| $D$ | $D$ | Diazotrophs | cells |
| $A$ | $A$ | $NO_3^-$-assimilators | cells |
| $M_{total}$ | $M_{total}$ | Mo available to D and A | mmol Mo |
| $M_D$ | - | Mo acquired by D | mmol Mo |
| $M_A$ | - | Mo acquired by A | mmol Mo |
| $Q_D$ | $Q_D$ | D's amount of Mo per a cell | mmol Mo $cell^{-1}$ |
| $Q_A$ | $Q_A$ | A's amount of Mo per a cell | mmol Mo $cell^{-1}$ |
| $\theta_D$ | $\theta_D$ | D's Mo requirement per a cell | mmol Mo $cell^{-1}$ |
| $\theta_A$ | $\theta_A$ | A's Mo requirement per a cell | mmol Mo $cell^{-1}$ |
| $\phi$ | $\phi$ | Population turnover | $day^{-1}$ |
| $r_D$ | $r_D$ | D's maximal growth rate | $day^{-1}$ |
| $r_A$ | $r_A$ | A's maximal growth rate | $day^{-1}$ |
| $k$ | - | D's competitive effectiveness in Mo uptake | - |
| - | $\mu_D$ | D's growth rate as a function of internal Mo | $day^{-1}$ |
| - | $\mu_A$ | A's growth rate as a function of internal Mo | $day^{-1}$ |
| - | $p_D$ | D's uptake rate of Mo | mmol Mo $cell^{-1}$ $day^{-1}$ |
| - | $p_A$ | A's uptake rate of Mo | mmol Mo $cell^{-1}$ $day^{-1}$ |
| - | $p_{max,D}$ | D's max uptake rate of Mo | mmol Mo $cell^{-1}$ $day^{-1}$ |
| - | $p_{max,A}$ | A's max uptake rate of Mo | mmol Mo $cell^{-1}$ $day^{-1}$ |
| - | $K_{S,D}$ | D's Mo uptake rate half maximum | mmol Mo |
| - | $K_{S,A}$ | A's Mo uptake rate half maximum | mmol Mo |
| - | $f$ | A's competitive effectiveness in Mo uptake | - |

**Table S2.** Collection of empirical Mo requirements

| | Organism | Mo:P (mmol:mol) | Mo in media (nmol $L^{-1}$) | Reference |
|---|---|---|---|---|
| **Diazotroph Bacteria** | *Cyanothece sp.* | 0.600 | 100 | (4) |
| | *Trichodesmium erythraeum* IMS101 | 0.304 | 97 | (10) |
| | *Crocosphaera watsonii* WH8501 | 0.077 | 97 | (10) |
| | *Anabaena flosaquae* | 0.100 | 100 | (4) |
| | *Nostoc spp. PCC 7120* | 0.160 | 108 | (11) |
| | *Nostoc spp. CCMP 2511* | 0.119 | 100 | (11) |
| | *Azotobacter vinelandii* | 0.470 | 100 | (12) |
| | *Azotobacter chroococcum* | 0.490 | 100 | (12) |
| | Mean | 0.290 | | |
| | Median | 0.232 | | |
| | Min | 0.077 | | |
| | Max | 0.600 | | |
| | Standard Deviation | 0.206 | | |
| **$NO_3^-$-assimilating Algae** | *Cyanophora paradoxa* | 0.030 | 100 | (4) |
| | *Chlorarachnion globosum* | 0.020 | 100 | (4) |
| | *Chlorarachnion reptans* | 0.040 | 100 | (4) |
| | *Eutreptiella sp.* | 0.026 | 100 | (4) |
| | *Rhodella maculata* | 0.020 | 100 | (4) |
| | *Rhodosorus marinus* | 0.020 | 100 | (4) |
| | *Porphyridium aerugineum* | 0.020 | 100 | (4) |
| | *Porphyridium purpureum* | 0.010 | 100 | (4) |
| | *Ochromonas sp.* | 0.008 | 100 | (4) |
| | *Rhodomonas salina* | 0.050 | 100 | (4) |
| | Mean | 0.024 | | |
| | Median | 0.020 | | |
| | Min | 0.008 | | |
| | Max | 0.050 | | |
| | Standard Deviation | 0.013 | | |

**Table S3.** Normalization of (10) data from Mo:C (μmol:mol) to Mo:P (mmol:mol)

| | ***Trichodesmium erythraeum* IMS101** | ***Crocosphaera watsonii* WH8501** |
|---|---|---|
| **Mo:C (μmol:mol)** | 1.33 | 0.45 |
| | 1.04 | 0.88 |
| | 6.62 | 0.57 |
| | 2.87 | 0.86 |
| | 2.47 | 0.65 |
| | | 0.90 |
| | | 0.75 |
| | | 0.77 |
| **Average** | 2.866 | 0.729 |
| **C:P (mol)** | 106 | 106 |
| **Mo:P (μmol:mol)** | 303.796 | 77.248 |
| **Mo:P (mmol:mol)** | **0.304** | **0.077** |

**Table S4.** T-test comparison of Mo:P requirements between diazotrophs and algae

| | ***Diazotrophs*** | ***Algae*** |
|---|---|---|
| Mean | 0.29 | 0.0244 |
| Variance | 0.0423 | 0.0002 |
| Observations | 8 | 10 |
| Hypothesized Mean Difference | 0 | |
| df | 7 | |
| t Stat | 3.646 | |
| P(T<=t) one-tail | 0.004 | |
| t Critical one-tail | 1.895 | |
| P(T<=t) two-tail | 0.008 | |
| t Critical two-tail | 2.365 | |
